# A chromosome-level genome assembly of the cabbage aphid *Brevicoryne brassicae*

**DOI:** 10.1101/2024.05.16.594461

**Authors:** Jun Wu, Guomeng Li, Zhimou Lin, Yangzi Zhang, Wenyun Yu, Rong Hu, Shuai Zhan, Yazhou Chen

## Abstract

The cabbage aphid, *Brevicoryne brassicae*, is a major pest on Brassicaceae plants, and causes significant yield losses annually. However, lacking genomic resources hinders the progress in understanding this pest at the level of molecular biology. Here, a high-quality, chromosomal-level genome was assembled for *B. brassicae* based on PacBio HIFI long-read sequencing and Hi-C data. The final assembled size was 429.99 Mb with a scaffold N50 of 93.31 Mb. Importantly, 96.2% of the assembled sequences were anchored to eight chromosomes. The genome recovered 98.50% of BUSCO genes and 92.30% of CEGMA genes, supporting the high level of completeness. By combining high-coverage transcriptome data, a total of 22,671 protein-coding genes and 3,594 lncRNA genes were annotated. Preliminary comparative genomic analyses were focused on genes related to host colonisation, such as chemosensory- and detoxification-related genes, as well as those encoding putative protein effectors and cross-kingdom lncRNA *Ya*. In summary, our study presents a contiguous and complete genome for *B. brassicae* that will benefit our understanding of the biology of it and other aphids.

## Background & Summary

*Brevicoryne brassicae*, the cabbage aphid, is a notorious pest that specializes in plants of the Brassicaceae family, including crops like rapeseeds, cabbage, and broccoli. *B. brassicae* cause damage to the plants directly through sap sucking from phloem tissues as well as indirectly by transmitting several plant viruses, which collectively result in significant yield losses to many Brassicaceae crops worldwide. *B. brassicae* is a nonhost-alternating species, meaning the life cycle can be completed on the herbaceous plants that are known as secondary hosts for host-alternating aphids^1^. The life cycle includes a sexual generation and several asexual generations. In the winter, *B. brassicae* produces sexual forms and overwinters in the egg stage. In warm seasons and regions, the life cycle is simplified to parthenogenetic reproduction. The winged females emerged when the population became dense and the host quality declined. Alates migrate to the distanced crops and parthenogenetically produce wingless ones, leading to population expansion exponentially and the escalation of aphid damage in the fields^2^.

Since the first aphid genome, the genome of *Acyrthosiphon pisum*, was published in 2010^3^, now dozens of aphid genomes are available, including important agricultural pests such as *Myzus persicae*^4,5^, *Aphis gossypii*^6^, *Diuraphis noxia*^7^, and valued recourse insects like *Schlechtendalia chinensis*^8^. The genomes have facilitated research on these aphids, leading to an improved understanding of molecular mechanisms of aphid biology. In contrast, studies on *B. brassicae* are much few, largely due to lacking genomic resources. Therefore, we constructed a high-quality *B. brassicae* genome at the chromosomal level using PacBio HIFI long reads and high-throughput chromosomal conformation capture (Hi-C) data. We annotated the genome for protein-coding and long non-coding RNA (lncRNA) genes, and performed phylogenic and evolutionary analysis with different aphid genomes. Our efforts will offer substantial support for a deeper understanding of *B. brassicae* and future studies into aphids.

After quality control filtering, we obtained a total of 29.00 Gb (∼67.44× depth) of PacBio long reads and 42.78 Gb (∼99.49× depth) of Illumina short reads. These reads were assembled into 131 contigs with an N50 length of 16.79 Mb (Table 1). Chromosome scaffolding based on Hi-C data resulted in eight chromosomes that contain 96.2% of scaffold sequences (Table 1). The chromosome number was confirmed by karyotype analysis (Fig. 1b). The final assembled genome is eight chromosomes with a total size of 429.99 Mb (Fig. 1c, Table 1). Chromosome lengths ranged from 16.74 Mb to 125.10 Mb (Fig.1d). The genome assembly is accurate at the gene level, containing 98.50% of arthropod BUSCO (Benchmarking Universal Single-Copy Ortholog) genes and 92.30% of CEGMA (Core Eukaryotic Genes Mapping Approach) genes (Table 1). Altogether, the assembly of *B. brassicae* genome are contiguous, accurate, and complete.

**Table 1.** Statistics for the chromosomal-level genome of *B. brassicae*.

| Statistical feature | Corresponding value |
| --- | --- |
| Genome size (Mb) | 429.99 |
| Number of chromosomes | 8 |
| Contig number | 131 |
| Contig N50 (Mb) | 16.79 |
| Scaffold number | 92 |
| Scaffold N50 (Mb) | 93.31 |
| BUSCO complete rate of the gnomes (%) | 98.50 |
| Percentage of CEGMA genes (%) | 92.30 |
| GC content (%) | 29.22 |
| All SNP | 203,407 |
| Percentage of SNPs (%) | 0.047 |
| Percentage of Heterozygosity SNPs (%) | 0.047 |
| Percentage of Homology SNPs (%) | 0.000010 |
| Protein-coding genes | 22,671 |
| lncRNA genes | 3,594 |
| Repeat (%) | 32.48 |

**Figure 1.**
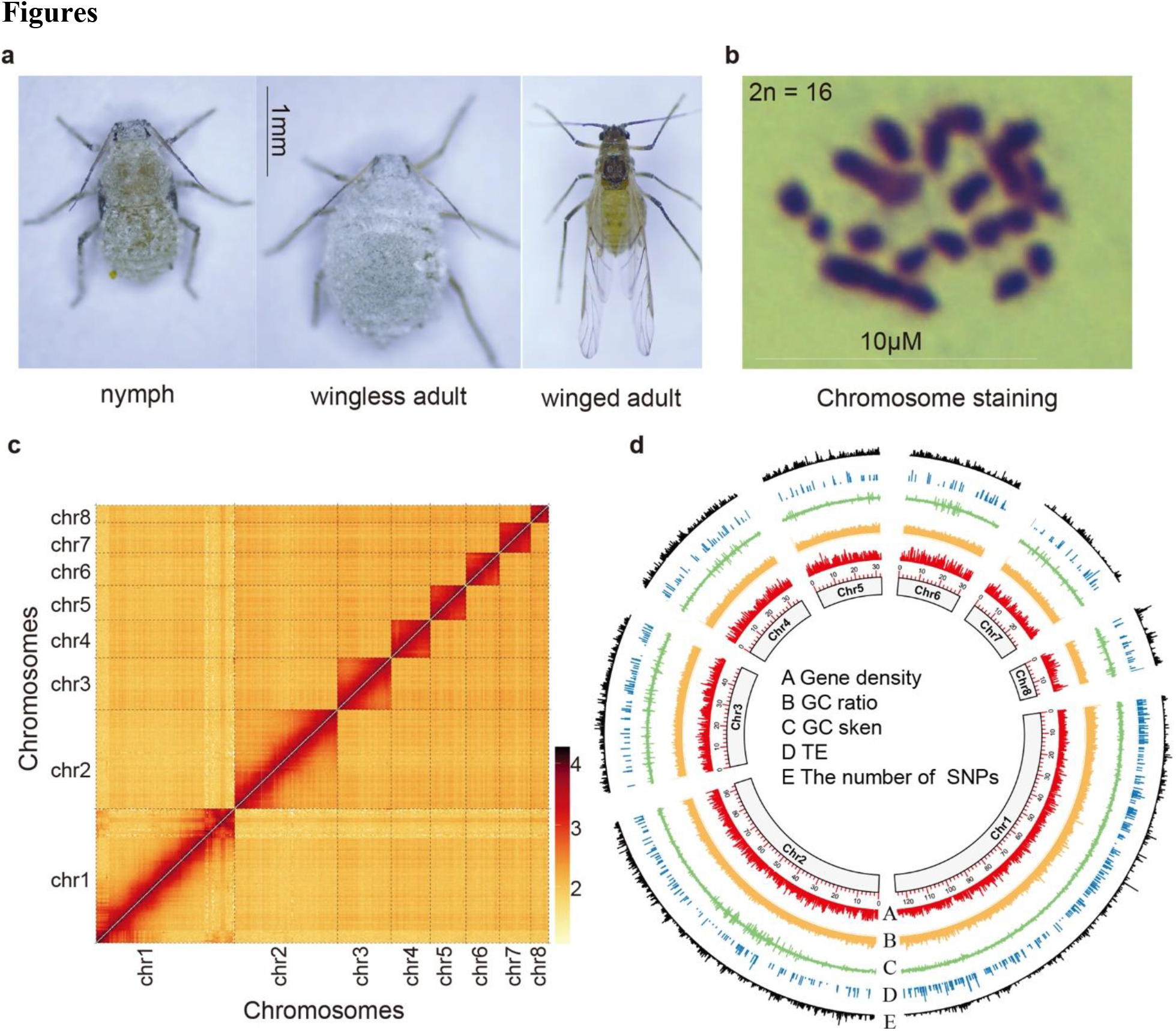
Morphological and genomic characteristics of *B. brassicae*. (**a**) Pictures of parthenogenesis *B. brassicae*: nymph, asexual female adult and alate adult. Bar = 1 mm. (**b**) Karyotype analysis of *B. brassicae*. Chromosomes (purple) were stained with Gurr’s Giemsa R66 (Giemsa), the diploidic *B. brassicae* has 16 chromosomes (2n (?) = 16). Bar = 10 μm. (**c**) Genome-wide contact matrix of *B. brassicae* generated using Hi-C data. The colour bar indicates the intensity of Hi-C interaction. Yellow indicates low, and red is high. (**d**) Circos plot overview of features of *B. brassicae* genome. Rings from inside to outside (A-E) represents gene density (A), GC ratios (B), GC skew (C), transposable element (D), and the number of SNPs (E).

Phylogenetic analysis revealed that *B. brassicae* diverged from *D. noxia* approximately 49.9 million years ago (MYA) and from other Macrosiphini species about 53.9 MYA (Fig. 2a). The *B. brassicae* genome showed high chromosomal synteny with *M. persicae*^5^ and *A. pisum*^9^ (Fig. 2b). Chromosome 1, the longest one (125.10 Mb) in the *B. brassicae* genome, showed massive synteny to the X chromosome of *M. persicae* and *A. pisum* and was therefore identified as X chromosome (Fig. 2b). Autosomes of *B. brassicae* genome have undergone extensive structural changes with many rearrangements between chromosomes, implying that aphid diversification is associated with dynamic changes in autosome structure.

**Figure 2.**
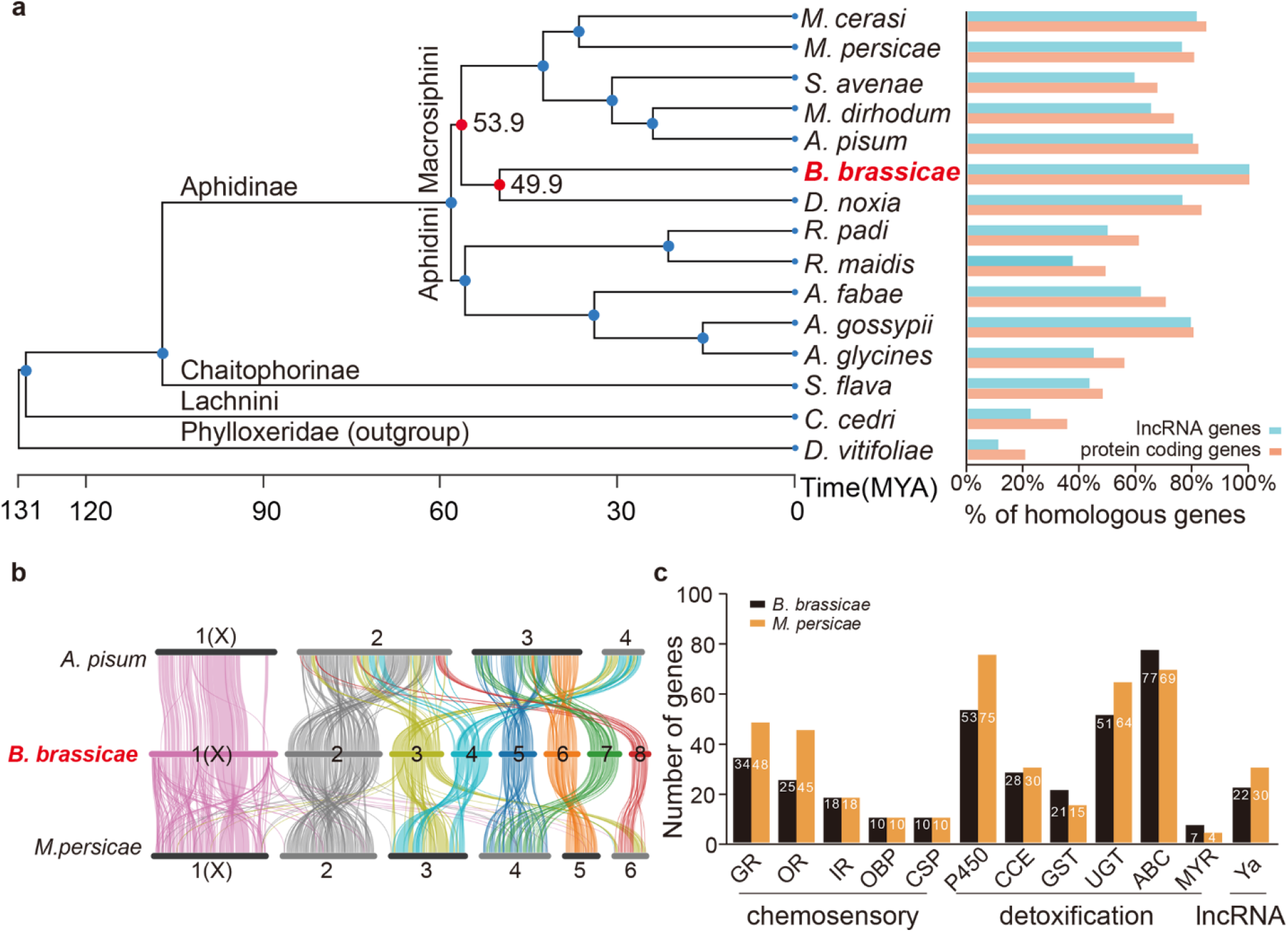
Comparative analysis of *B. brassicae* genome. (**a**) Phylogenetic tree constructed based on the 484 single-copy genes of 15 aphids (*Myzus cerasi, Myzus persicae, Sitobin avenae, Metopolophium dirhodum, Acyrthosiphon pisum, B. brassicae, Diuraphis noxia, Rhopalosiphum padi, Rhopalosiphum maidis, Aphis fabae, Aphis gossypii, Aphis glycines, Sipha flava, Cinara cedri*, and *Daktulosphaira vitifoliae*). The bar graph displays the proportion of homology of randomly selected protein-coding and lncRNA genes (600 of each) of *B. brassicae*. (**b**) Synteny of the genomes of *A. pisum, B. brassicae* and *M. persicae*. (**c**) The number of genes related to host adaptation in *B. brassicae* (specialist) and *M. persicae* (generalist). Chemosensory-related gene families, including gustatory receptors (GR), olfactory receptors (OR), ionotropic receptors (IR), odorant-binding proteins (OBP), and chemosensory proteins (CSP); detoxification gene families, including cytochrome P450 (P450), carboxylesterases (CCE), glutathione S-transferases (GST), UDP-glucuronosyltransferases (UGT), ATP-binding cassette transporters (ABC), and myrosinase (MYR) genes, as well as lncRNA *Ya* gene family.

Using chromosome-level genome assemblies, we annotated protein-coding genes and lncRNA genes using evidence from 90.31 Gb of RNA sequencing (RNA-seq) data (63.60 Gb un-stranded and 26.71 Gb stranded). In total, 22,671 protein-coding genes and 3,594 lncRNA genes were annotated (Table 1). Conservation of protein-coding sequences is high among different aphid species, and lncRNA sequences trended to be more divergent (Fig. 2a), suggesting rapid turn-over of lncRNA genes during the speciation. Additionally, we annotated 154 loci of miRNAs, 332 loci of tRNA, 837 loci of rRNAs, and 124 loci of snRNAs (Table 3). We also identified 141.19 Mb of repeating sequences accounting for 32.48% of the genome assembly (Table 2).

**Table 2.** Statistics of the transposable elements in the genome of *B. brassicae*.

| Type | Denovo+Rebase |  | TE Proteins |  | Combined TEs |  |
| --- | --- | --- | --- | --- | --- | --- |
|  | Length<br>(bp) | % in<br>genome | Length<br>(bp) | % in<br>genome | Length<br>(bp) | % in<br>genome |
| DNA | 36,542,400 | 8.50 | 3,241,425 | 0.75 | 37,903,104 | 8.81 |
| LINE | 5,782,148 | 1.34 | 1,460,641 | 0.34 | 6,485,087 | 1.51 |
| SINE | 3,697 | 0.00 | 0.00 | 0.00 | 3,697 | 0.00 |
| LTR | 74,665,259 | 17.36 | 4,788,589 | 1.11 | 75,112,021 | 17.47 |
| Unknown | 28,483,967 | 6.62 | 0.00 | 0.00 | 28,483,967 | 6.62 |
| Total | 140,326,512 | 32.63 | 9,490,171 | 2.21 | 141,191,380 | 32.84 |

**Table 3.** Genomic annotation of miRNA, tRNA, rRNA, and snRNA loci in the genome of *B. brassicae*.

|  | Type | Copy number | Average length (bp) | Total length (bp) | Coverage of genome (%) |
| --- | --- | --- | --- | --- | --- |
|  | miRNA | 154 | 106.91 | 16,464 | 0.0038 |
|  | tRNA | 332 | 75.01 | 24,902 | 0.0058 |
|  | **rRNA** | **837** | **190.92** | **159,798** | **0.037** |
| **rRNA** | 18S | 260 | 258.65 | 67,248 | 0.016 |
|  | 28S | 574 | 160.88 | 92,343 | 0.021 |
|  | 5.8S | 0 | 0 | 0 | 0.00 |
|  | 5S | 3 | 69 | 207 | 0.000048 |
|  | **snRNA** | **124** | **136.21** | **16,890** | **0.0039** |
| **snRNA** | CD-box | 44 | 108 | 4,752 | 0.0011 |
|  | HACA-box | 0 | 0 | 0 | 0.00 |
|  | splicing | 70 | 148.89 | 10,422 | 0.0024 |
|  | scaRNA | 10 | 171.6 | 1,716 | 0.00040 |
|  | Unknown | 0 | 0 | 0 | 0.00 |

*B. brassicae* is a specialist aphid that is restricted to the Brassicaceae plants^10^. To successfully colonize the host plants, *B. brassicae* has evolved strategies to recognize host plants and overcome plant defence^11–13^. Thus, we annotated chemosensory-related genes for gustatory receptors (GRs), odorant receptors (ORs), ionotropic receptors (IRs), odorant-binding proteins (OBPs) as well as chemosensory proteins (CSPs). The numbers of genes encoded for GRs, ORs, IRs, OBPs and CSPs in *B. brassicae* genome were 34, 25, 18, 10, and 10, respectively. The number of GR and OR genes were less than that in the generalist aphid *M. persicae* (Fig. 2c). In addition, we annotated detoxification-related genes and identified 53 genes for cytochrome P450 (P450), 28 for carboxyl/choline esterase (CCE), 21 for glutathione-S-transferase (GST), 51 for UDP-Glycosyltransferase (UGT), 77 for ATP-binding cassette transporters (ABC), and 7 for myrosinases (MYR) (Fig. 2c). Compared to *M. persicae* genome, *B. brassicae* genome encodes fewer genes for P450, CCE, and UGT, but more genes for GST, ABC and MYR. These results suggested concerted expansion and contraction of chemosensory-related and detoxification-related genes in *B. brassicae* genomes that contributed to its specialist lifestyle.

During feeding, aphids often secrete molecular weapons, such as effector proteins^14^ and cross-kingdom lncRNAs^15^, to repress plant defences and promote aphid colonization. In the *B. brassicae* genome, we identified 1,015 putative protein effectors, of which 980 were conserved among many aphid species and 35 were specific to the *B. brassicae* (Fig. 2a). In addition to protein effectors, *M. persicae* lncRNA *Ya* are translocated into host plants and act as virulence factors^15^. Here, we identified 22 *Ya* genes in *B. brassicae* genome. Compared to 30 *Ya* genes in *M. persciae, Ya* genes in *B. brassicae* are fewer, suggesting *Ya* family may be associated with the host breadth of aphids.

This study presents the high-quality chromosome-level genome assembly of *B. brassicae* and comprehensive annotations, which provide an invaluable genomic resource for understanding the genetic, evolutionary, and ecological issues of the cabbage aphid, and further offer the possibility to implement integrated pest management of this pest.

## Methods

### Sample collection and genome sequencing

The *B. brassicae* larvae were collected from rapeseed fields in Lanzhou, Gansu province, China. The insects were reared on the rapeseed plants in the lab. Genomic DNA was extracted from adult insects using the CTAB (cetyltrimethylammonium bromide) method^16^ for Illumina, PacBio, and Hi-C sequencing. The quality of the genomic DNA was validated using NanoDrop 2000C spectrophotometer and 1.5% agarose gel electrophoresis. Briefly, the genomic DNA sample was fragmented by sonication to a size of 350 bp. Then DNA fragments were end-polished, A-tailed, and ligated with the full-length adapter using SMRTbell Express Template Prep Kit 2.0 (Pacific Biosciences, Menlo Park, CA, USA) for Illumina sequencing. Libraries with a 350 bp inserted fragment were constructed and sequenced on the Illumina NovaSeq 6000 platform. After removing adapter sequences and low-quality reads, 42.78 Gb of clean data were obtained for subsequent analysis.

For PacBio HiFi sequencing, genomic DNA was sheared into ∼15 kb fragments using g-Tubes (Covaris, Woburn, MA, USA) and purified using 0.45 × AMPure PB beads (Beckman Coulter, Brea, CA, USA) to construct SMRT bell libraries. Size selection was performed using BluePippin (Sage Science, Beverly, MA, USA) to collect 15-18 kb fragments. After annealing the primers and binding Sequel DNA polymerase to SMRT bell templates, sequencing was performed using one SMRT cell 1 M on the Sequel System (Biomarker Technologies). Finally, a total of 29.00 Gb of subreads were obtained, with an average read length of 10.45 kb, resulting in 67.44×coverage of the *B. brassicae* genome. To achieve chromosome-level assembly, the Hi-C technique was used to identify contacts between different regions of chromatin filaments. The Hi-C library was constructed following the standard library preparation protocol and sequenced on the Illumina NovaSeq 6000 platform, and 60.78 Gb of 150-bp paired-end clean reads were obtained.

### RNA extraction and transcriptome sequencing

30 first-instar nymphs, 15 asexual adults, and 15 winged female adults collected from rapeseed plants in the lab were used for RNA extraction. Total RNA was extracted using TRIzol reagent (Invitrogen, Carlsbad, CA, USA) following the manufacturer’s instructions. The concentration of the isolated RNA was measured using a NanoDrop 2000C spectrophotometer (Thermo Fisher Scientific, Pittsburgh, PA, USA). RNA quality was evaluated using 1.5% agarose gel electrophoresis. RNA integrity was quantified using an Agilent 5400 Fragment Analyzer (Agilent, Santa Clara, CA, USA). RNA-seq libraries were constructed using the NEBNext® Ultra™ RNA Library Prep Kit (NEB, Ipswich, MA, USA) following the manufacturer’s instructions. Libraries were then sequenced on the Illumina NovaSeq 6000 platform, and 63.60 Gb un-stranded and 26.71 Gb stranded 150-bp paired-end reads were obtained and used for gene prediction.

### Genome estimation and assembly

PacBio HIFI long-read data were used to generate a contig-level assembly of the *B. brassicae* genome. A preliminary assembly was generated using WTDBG2 v2.5^17^ with the default parameters. After correcting for short-read using Pilon v1.23^18^, the *B. brassicae* genome assembly was generated, which consisted of 131 contigs with a total length of 429.99 Mb and a contig N50 of 16.79 Mb. After removing the low-quality reads and adaptor sequences, 29.00 Gb of clean data were generated from the Hi-C library and mapped to the draft *B. brassicae* genome using BWA v0.7.10^19^ with the default parameters. Uniquely aligned read pairs were further processed using HiC-Pro v2.10.0^20^ to assess and eliminate invalid read pairs, including dangling ends, re-ligation, self-cycle, and dumped pairs. Valid interaction pairs for scaffold correction were used to cluster, order, and orient the contigs onto chromosomes using LACHESIS v2e27abb^21^ with the default parameters. Ultimately, eight chromosomes with a scaffold N50 of 93.31 Mb were constructed, covering a span of 429.99 Mb and representing 96.19% of the draft genome assembly.

### Genomic repeat annotation

Repeat sequences mainly include tandem and interspersed repeats, the latter being primarily transposable elements (TEs). The repeat TE sequences were annotated using a combination of homology-based and *de novo* approaches. We initially customized a *de novo* repeat library using RepeatModeler v2.0.2a^22^, LTR_FINDER^23^, and RepeatScout^24^ based on assembly sequences with default parameters. The predicted repeats were subsequently classified using the PASTE Classifier v1.0^25^, and the results were combined with the database of Dfam v3.2^26^ to construct a species-specific TE library without redundancy. The TE sequences were identified by homology searching against the library using RepeatMasker v4.10^27^ and Repeatproteinmask^27^. Ultimately, 141.19 Mb of TE sequences were identified, accounting for 32.84% of the genome assembly. Long terminal repeats (LTR) were the largest category of transposable elements, representing 17.47% of the genome, followed by DNA transposons, representing 8.81% of the genome, unknown repeated sequences and long interspersed nuclear elements (LINE), accounting for 6.62% and 1.51% of the whole genome (Table 2).

### Protein-coding gene annotation

An integrated approach based on *B. brassicae* transcriptome and protein homologs from other aphids was used for predicting protein-coding genes on the reference genome being masked with repeats. The RNA-seq data from pooled wingless/winged asexual females and nymphs was used. Reads were mapped to the reference genome using HISAT2 v2.2.1^28^ with default parameters and processed by SAMtools v0.1.18^29^. The alignment results were provided for Braker1^30^, which generated a transcriptome-based gene set. Furthermore, protein-coding genes of related species, including *M. persicae, A. pisum*, and *A. gossypii*, and the model insect *Drosophila melanogaster*, were filtered with isoforms and provided for Braker2^31^, which generated another homolog-based gene set. The two independent gene sets were compared at both exon and transcript levels to generate a consensus gene set. To do this, unique models non-overlapped to each other were selected first, while the models with the disparity between the two approaches were further checked based on evidence of homolog alignment and transcriptome to reserve the best one.

### Phylogenetic tree construction and genome synteny analyses

We identified 484 single-copy genes using OrthoFinder v2.5.4^32^ based on protein sequences from 15 aphid genomes, including 7 Macrosiphini species (*M. cerasi*^33^, *M. perisicae*^5^, *Sitobion avenae*^34^, *Metopolophium dirhodum*^35^, *A. pisum*^9^, *B. brassicae*, and *D. noxia*^7^), 5 Aphidini species (*Rhopalosiphum maidis*^36^, *Rhopalosiphum padi*^33^, *Aphis fabae*^33^, *A. gossypii*^6^, *Aphis glycines*^33^), one species from each subfamily of Chaitophorinae (*Sipha flava*^37^), Lachnini (*Cinara cedri*^38^), Phylloxeridae (*Daktulosphaira vitifoliae*^39^). The protein sequences of the single-copy genes were concatenated and aligned automatically by OrthoFinder and generated a multiple sequence alignment file, which was used for phylogenetic analysis. For the phylogenetic tree reconstruction, ProTest v3.2^40^ was used first and found “JTT + I + G4” to be the best model, which was later used in the maximum likelihood phylogenetic tree reconstruction using RAxML v8.2.12^41^. We used iTOL v6^42^ for tree visualization.

For the genome synteny analysis, the 1:1:1 orthologs among *B. brassicae, M. perisicae*, and *A. pisum* genomes were extracted from OrthoFinder’s result and fed to MCScanX_h^43^, which was used with “-b 2” option to get the inter-species collinearity among *B. brassicae, M. perisicae*, and *A. pisum*. SynVisio^44^ was used to visualize the genome synteny.

### Identification of putative aphid effector proteins

Putative proteinic effectors are characterized as secreted proteins with no transmembrane properties. Protein sequences of 22,671 genes were analyzed separately by SignalP v6.0^45^ to identify secretion signals and by TMHMM v2.0^46^ to identify transmembrane domains, which led to the identification of 1,015 putative protein effectors.

### Annotation of long non-coding RNAs

The process of identifying lncRNA genes in the *B. brassicae* genome is divided into three main steps. Firstly, mapping of stranded RNA-seq to the *B. brassicae* genome. The raw reads are subjected to quality control using Fastp v0.23.4^47^ with default parameters to ensure data integrity for downstream analyses. The processed reads are aligned to *B. brassicae* genome using HISAT2 v2.2.1^28^. The aligned reads are assemleid into transcripts by StringTie v2.2.1^48^. The Gffread v0.12.7^49^ with the parameters “-V -H -U -N -P -J -M -K -Q -Y -Z -F --keep-exon-attrs” was used for extracting the assembled transcript sequences. Secondly, LGC (Long Genomic Region Classifier) v1.0^50^ is used for identifying transcripts with non-coding features based on the relationship between ORF (open reading frame) Length and GC content. Meanwhile, assembled transcripts were subjected to the CPC2 (Coding Potential Calculator 2)^51^ to calculate the coding potential. The intersection of the results from LGC and CPC2 were the putative lncRNAs. Thirdly, the putative lncRNA transcripts were screened by rFAM v14.3^52^ to eliminate housekeeping RNAs, such as rRNA, tRNA, and snoRNA.

To annotate the *Ya* gene family in the *B. brassicae* genome, the sequence of *M. persicae Ya1* (*MpYa1*) was used as a query. BLASTn^53^ was utilized to perform sequence alignment of the *MpYa1* sequence against the annotated lncRNA transcripts with an E-value cutoff of less than 10^-5 and a 30% similarity threshold. The final alignment resulted in the identification of 22 *Ya* genes in the *B. brassicae*.

### Gene family identification

To annotate the detoxification- and chemosensory-related genes of *B. brassicae*, amino acid sequences of those genes reported in other aphid species were used as the query in the Diamond blast v0.8.29^54^ to identify homologies with E-value less than 10^-5. The sequences of the detoxification-related genes, including cytochrome P450 (P450), carboxylesterases (CCE), glutathione S-transferases (GST), UDP-glucuronosyltransferases (UGT), ATP-binding cassette transporters (ABC), and myrosinase (MYR) genes, were downloaded from the InsectBase v2.0^55^. The sequences of the chemosensory-related genes, including gustatory receptors (GRs), olfactory receptors (ORs), ionotropic receptors (IRs), odorant-binding proteins (OBPs), and chemosensory proteins (CSPs) were obtained from published papers^56,57^.

### Karyotype analysis

The number of chromosomes was confirmed by karyotype analysis, indicating that diploidic *B. brassicae* has 16 chromosomes (2n = 16). The Gurr’s Giemsa R66 chromosome staining method^58^ was used. Briefly, chromosome squash preparations are made from young embryos dissected from parthenogenetic adult aphids. The embryos were treated in 0.75% of potassium chloride, and then fixed in a freshly prepared mixture of absolute methanol and glacial acetic acid (3:1 in volume) for 10 minutes. Next, the embryos were carefully transferred onto a pin’s tip, subsequently were moved to a clean microscope slide with a small drop of 45% propionic acid (5 minutes), squashed with a coverslip then dried for 24 hours at room temperature. HCl solution (0.2 M) is applied dropwise for 30 minutes at room temperature, followed by rinsing with distilled water and immersion in a 5% saturated Ba(OH)_2_ solution. The sample is then treated in a 60°C constant temperature water bath for 3 minutes. After the treatment, it is briefly processed in HCl solution (0.2 M) to interrupt the reaction with the strong base, rinsed with distilled water, and air-dried at room temperature. Subsequently, the sample is stained with 5% Giemsa stain (pH 7.0) for 30 minutes, air-dried at room temperature, and examined and photographed under an optical microscope.

### Data Records

The genome sequencing data (PacBio, Illumina and HiC) of *B. brassicae* have been submitted to the Sequence Read Archive (SRA) at the National Center for Biotechnology Information (NCBI) under the BioProject of PRJNA1099426^59–61^. The RNA sequencing (RNA-seq) data generated in this study have been deposited in the SRA at the NCBI under the BioProject accession number PRJNA1104693 and this submission includes a total of 9 un-stranded RNA-seq data^62–70^ and 3 stranded RNA-seq data^71–73^. The *B. brassicae* genome assembly FASTA and GFF files, the annotation GTF files of protein-coding genes, the annotation files including PFAM, KEGG and GO, the annotation files of several regulatory elements including transposable element, lncRNA and miRNA, the annotation files of tRNA, rRNA, and snRNA loci, and the protein sequences of detoxification- and chemosensory-related genes have been deposited in the Figshare database^74^.

### Technical Validation

#### Assessing the validity of gene prediction and annotation

The number of chromosomes was confirmed by karyotype analysis, indicating that diploidic *B. brassicae* has 16 chromosomes (2n = 16). The complete BUSCOs under genome mode were used to assess the genome completeness. A total of 98.50% complete BUSCOs were identified, including 95.85% single-copy BUSCOs, 2.57% duplicated BUSCOs, 0.20% fragmented BUSCOs, and 1.38% missing BUSCOs. 95.16% of completeness of CEGs was identified based on 248 ultra-conserved CEGs.

## Code availability

No specific script was used in this work. All commands and pipelines used in data processing were executed according to the manual and protocols of the corresponding bioinformatic software.

## Acknowledgements

This project is funded by the National Key Research and Development Program of China (project No. 2023YFF1000700), and supported by Hubei Hongshan Laboratory (project No. 2022hszd026 to YC), the Startup Foundation for Advanced Talents at HZAU to YC, the Fundamental Research Funds for the Central Universities (Program No. 2022ZKPY003 to YC), and the Wuhan Yingcai Talent Program to YC.

## Author contributions

Y.C. conceived and led the research, Y.C., G.L., J.W., Z. L and Y.Z. were involved in sample collection, preparation and genome assembly. S.Z., Y.C., J.W. and G.L. contributed to gene prediction and annotation, data analysis. W.Y conducted the karyotype analysis. Y.C. and R.H. contributed to data management. Y.C. and J.W. wrote the manuscript and all authors read, revised and approved the final version of the manuscript. Y.C. supervised the project.

## Additional information

**Correspondence** and requests for materials should be addressed to Y.C.

